# Within-Patient Evolution of *Pseudomonas aeruginosa* Populations During Antimicrobial Treatment

**DOI:** 10.1101/2025.09.19.677277

**Authors:** Giuseppe Fleres, Ellen G. Kline, Kevin M. Squires, Tyler Tate, Hannah M. Creager, Ryan K. Shields, Daria Van Tyne

**Affiliations:** Division of Infectious Diseases, University of Pittsburgh School of Medicine; Center for Evolutionary Biology and Medicine, University of Pittsburgh School of Medicine; Center for Innovative Antimicrobial Therapy, University of Pittsburgh School of Medicine; Antibiotic Management Program, University of Pittsburgh Medical Center

**Keywords:** culture-enriched metagenomics, *Pseudomonas aeruginosa*, within-host evolution

## Abstract

Multidrug-resistant (MDR) *Pseudomonas aeruginosa* infections pose a major challenge to effective treatment. Understanding genomic adaptations during antimicrobial therapy in patients infected with this pathogen is crucial for preventing therapeutic failure. Here we investigated the population diversity and evolution of *P. aeruginosa* collected longitudinally from six patients who evolved multidrug-resistant infections. Serial *P. aeruginosa* clinical isolates (n=63) and culture-enriched metagenomic population samples (n=39) were collected and subjected to whole-genome sequencing. The resulting data were used to characterize and compare the species composition, multi-locus sequence types (STs), and resistance-associated mutations present within each sample type. Single-isolate sequencing showed that each patient was infected with a single *P. aeruginosa* strain that accumulated mutations and became increasingly more resistant over time. Mutations in genes associated with beta-lactam resistance, including *ampC, ftsI*, and *mexR*, arose over time and corresponded with changes in antimicrobial susceptibility in single isolates. Species profiling of culture-enriched metagenomic populations revealed that all samples contained *P. aeruginosa*, but also additional Gram-negative pathogens. Metagenomic analysis of culture-enriched populations identified resistance-associated mutations at low frequency, many of which were not identified in single isolates from the same sample. In some cases, resistance-associated mutations initially detected at low frequency rose to fixation after antimicrobial treatment. Overall, this study shows that population-based metagenomic sequencing effectively captures within-patient genomic diversity of *P. aeruginosa* during antimicrobial therapy, and could aid the detection and interpretation of resistance-associated mutations in this pathogen.

**Importance:** *Pseudomonas aeruginosa* infections are notoriously difficult to treat and are associated with high rates of morbidity and mortality. While the genetic basis of resistance in *P. aeruginosa* is well documented *in vitro*, less is known about how resistance evolves within patients during antibiotic therapy. Standard approaches based on analysis of clonal isolates may miss within-patient diversity, potentially overlooking low-frequency mutations that contribute to treatment failure. In this study, we integrated single-colony whole-genome sequencing with culture-enriched metagenomic sequencing to monitor the evolution of *P. aeruginosa* populations in patients receiving antibiotic therapy. This approach enabled the detection of emerging resistance mutations, such as low-frequency variants in *ampC* and *ftsI*, before these variants rose to fixation. It also revealed genetically resistant subpopulations missed by isolate sequencing alone. Together, our findings highlight the value of population-based metagenomic sequencing in capturing bacterial adaptation during infection, and underscore its potential to improve resistance surveillance and guide personalized antimicrobial therapy.

## Observation

*Pseudomonas aeruginosa* is recognized as a serious public health threat by the Centers for Disease Control and Prevention and has been classified by the World Health Organization as a critical priority pathogen, emphasizing the urgent need for new antimicrobial therapies^1^. A variety of intrinsic resistance mechanisms, combined with a remarkable capacity to acquire additional resistance determinants, make *P. aeruginosa* infections increasingly difficult to treat^2^. Infections caused by multidrug-resistant (MDR) strains are particularly concerning and often necessitate the use of newer β-lactam/β-lactamase inhibitor combinations or siderophore cephalosporins to manage^3,4^. However, the efficacy of these agents is frequently undermined by the emergence of diverse and often convergent resistance mechanisms. These include upregulation of multidrug efflux systems (e.g., MexAB-OprM), inactivation or loss of porins (OprD), mutations affecting chromosomal β-lactamase expression (AmpC), and point mutations in essential antibiotic target genes like *ftsI* (encoding penicillin-binding protein 3), *gyrA*, and *gyrB*^4,5^. Alterations in regulatory genes such as *lasR* further contribute to phenotypic heterogeneity by modulating quorum sensing pathways, which can affect antibiotic tolerance and persistence^2^. These adaptive mutations frequently arise under selective therapeutic pressures, facilitating rapid within-host evolution and complicating effective treatment^5^.

Although resistance mechanisms in *P. aeruginosa* have been well characterized *in vitro* and among serial clinical isolates collected from infected patients^6,7^, much less is known about how infecting *P. aeruginosa* populations evolve within patients during antibiotic therapy^8^. Treatment decisions are typically guided by antimicrobial susceptibility testing and whole-genome sequencing analysis of a single bacterial clone, assuming that it reflects the entire infecting population. However, this approach offers limited insight into within-host diversity and often misses low-frequency variants that may influence treatment outcomes^9,10^. Culture-enriched metagenomic sequencing addresses these limitations by pooling all colonies from a selective culture plate and performing shotgun metagenomic sequencing, enabling a more comprehensive analysis of population-level diversity and evolution during infection^9,11^.

Here we investigated the within-host evolution of *P. aeruginosa* populations in six patients who developed MDR infections during treatment and compared results obtained from clonal and culture-enriched metagenomic sequencing. We hypothesized that reliance on single-colony testing would underestimate the genotypic and phenotypic diversity of *P. aeruginosa* within the infected host, particularly under antimicrobial selection. A total of 63 clinical isolates and 39 culture-enriched metagenomic populations were collected from six patients, sequenced, and analyzed (Figure 1) (Supplemental File 1) (Supplemental Methods). In all patients, susceptibility profiles of *P. aeruginosa* isolates progressively shifted toward increasing resistance over the course of treatment. This aligns with prior studies documenting the rapid evolution of resistance in *P. aeruginosa* under antibiotic stress, especially during prolonged therapy in chronic infections^12,13^. In two patients (Patients 1 and 6), isolates collected at the final time point exhibited a more susceptible phenotype compared to earlier isolates, suggesting that resistant bacteria may have been outcompeted or replaced, potentially due to changes in treatment, host immunity, or fitness costs associated with resistance^14^.

**Figure 1.**
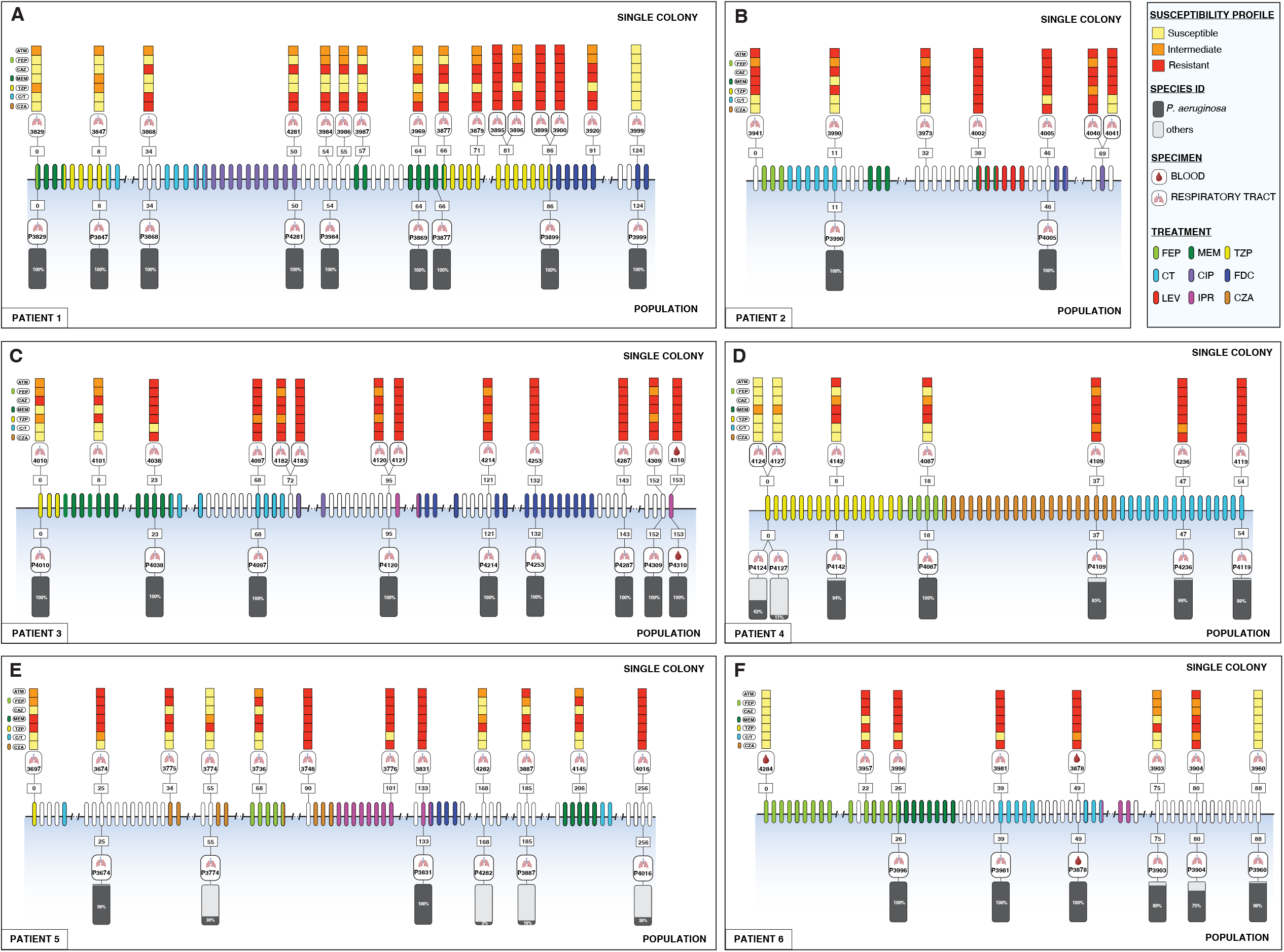
Timeline of *P. aeruginosa* infection and treatment for each patient. The panel shows longitudinal sampling of *P. aeruginosa* single colony isolates and culture-enriched metagenomic populations during recurrent infections for each patient. The upper section of each panel depicts single colony isolates, while the lower section shows corresponding population samples. Colored squares indicate the antimicrobial susceptibility profile of each isolate, and grey bars show the proportion of each population composed of *P. aeruginosa* or other bacterial species. β-lactam antibiotics tested included aztreonam (ATM), cefepime (FEP), ceftazidime (CAZ), meropenem (MEM), piperacillin/tazobactam (TZP), ceftolozane/tazobactam (C/T), and ceftazidime/avibactam (CZA). Ovals in each timeline are colored based on antimicrobial treatment administered to each patient, including FEP, MEM, TZP, C/T, ciprofloxacin (CIP), cefiderocol (FDC), levofloxacin (LEV), imipenem/relebactam (IPR), and CZA. Sample identifiers, source (respiratory tract or blood), and collection timepoints (in days) are indicated above and below the timeline.

Species profiling of culture-enriched metagenomic samples confirmed the presence of *P. aeruginosa* in all samples (Figure 1). Metagenomic samples from three patients (Patients 1–3) contained only *P. aeruginosa*, while the other three patients (Patients 4–6) exhibited polymicrobial infections including *P. aeruginosa* and other organisms (Supplemental Figure 1). In each case, the same species were identified as part of standard-of-care testing in the clinical microbiology laboratory, suggesting that this approach accurately captured polymicrobial infections.

Single-colony sequencing revealed a *P. aeruginosa* sequence type (ST) that was unique to each patient, and all colonies sequenced from the same patient belonged to the same ST (Supplemental Figure 2A). Analysis of within-patient single nucleotide polymorphisms (SNP) among *P. aeruginosa* isolates collected from the same patient revealed low genetic diversity in five of the six patients, with pairwise SNP differences ranging from 0 to 23. This narrow range suggested that each patient was infected with a single strain that underwent clonal expansion during infection. In contrast, one patient (Patient 1) displayed markedly higher within-host diversity, with SNP distances reaching up to 411 (Supplemental Table 2). This elevated level of genetic variation could be attributed to the presence of a hypermutator clone, which accelerates mutation rates and promotes rapid genetic diversification, particularly under selective pressures such as antibiotic treatment or the host immune response^15^. Supporting this interpretation, multiple mutations were identified in DNA mismatch repair genes from this patient’s isolates, including W181R and S324P in *mutS* and L35P in *mutL*, indicating a defective mismatch repair system.

We next explored whether culture-enriched metagenomic sequencing could enhance the detection of resistance-associated mutations that might be missed by single-colony sequencing. In our analysis of 11 resistance-associated genes across 6 patients, culture-enriched metagenomic sequencing provided additional insight in 19/66 (29%) cases. In 84% of these cases (n=16/19), this was due to the detection of resistance-associated mutations that were not identified by single-colony sequencing (Supplemental Table 3). Comparison of single-colony and culture-enriched metagenomic sequencing revealed distinct interpretations depending on the method used (Figure 2). In Patient 3, single-colony sequencing identified a low-frequency mutation in *ampC* (P243S) but failed to detect a co-occurring mutation (E247K) that was present at over 80% frequency in the population (Figure 2A). This highlights a key limitation of the single-colony approach, which might randomly capture minority variants while missing the dominant genotype, ultimately leading to an incomplete or misleading resistance profile. In both Patients 3 and 4, mutations in *ftsI* were identified at low frequency in population samples collected at earlier time points, and the same variants were later detected in single colonies as well as in the populations at high frequency (Figure 2A and 2B). This illustrates how population sequencing can detect emerging mutations before they become dominant. Finally, in Patient 6 we identified mutations at allele frequencies less than 50% that were exclusively detected in population samples and were never observed in single-colony genomes (Figure 2C). Together these findings emphasize the higher sensitivity of culture-enriched metagenomic sequencing for capturing within-host diversity.

**Figure 2.**
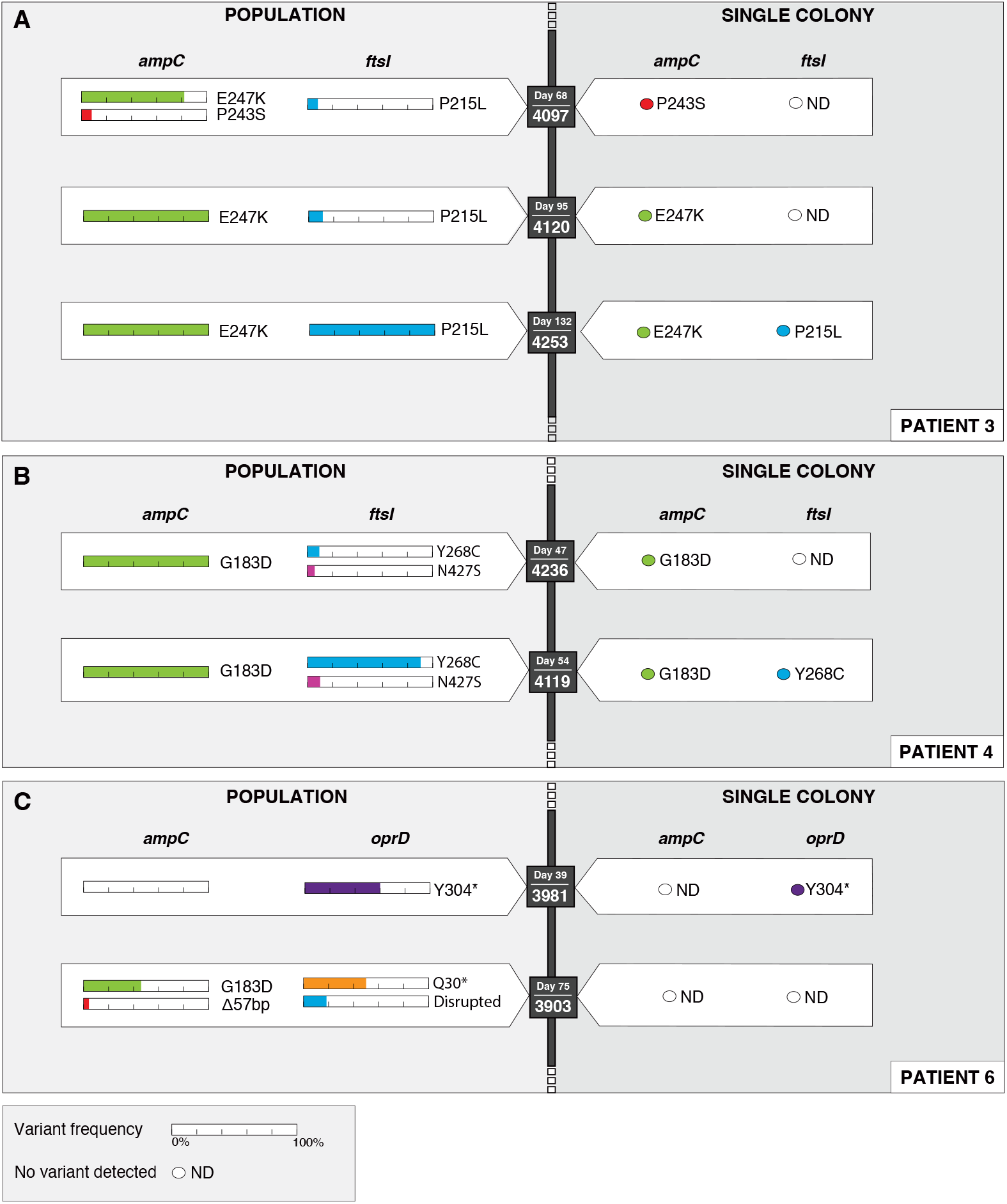
Comparison of resistance-associated mutations detected in culture-enriched metagenomic populations versus single colonies in three patients with *P. aeruginosa* infections. Each panel represents a subset of the isolate and population data collected from each patient, and displays mutations identified in population sequencing (left) and in single colony isolate whole-genome sequences (right) at corresponding sampling timepoints. Nonsynonymous mutations, nonsense mutations, and structural variants are shown in beta-lactam resistance-associated genes *ampC, oprD*, and *ftsI*. Colored bars represent the allele frequency (%) of each mutation in metagenomic populations. Sample ID and collection day are indicated next to each genotype.

Here we focused on a targeted set of genes known to be associated with antibiotic resistance. While this approach allowed for in-depth analysis of clinically relevant resistance mechanisms, it likely underrepresents the broader genomic heterogeneity within infecting *P. aeruginosa* populations. Other adaptive changes (e.g., mutations related to antibiotic tolerance, biofilm formation, metabolic remodeling, etc.) may also contribute to treatment failure but were not systematically assessed here. Future studies incorporating unbiased, genome-wide analyses could provide a more comprehensive view of within-host evolution during infection and treatment.

In summary, this study demonstrates that culture-enriched metagenomic sequencing provides enhanced resolution of within-host evolution in *P. aeruginosa* during antimicrobial treatment. Compared with prior studies, here we used a combined approach of single-colony and culture-enriched metagenomic sequencing to achieve higher-resolution tracking of within-host evolution during antibiotic treatment in infected patients. We found that population-level sequencing consistently identified low-frequency variants that were missed by single-colony sequencing, revealing a more comprehensive picture of the adaptive landscape in these patients. As WGS becomes more commonly used in clinical practice, culture-enriched metagenomic sequencing could provide greater sensitivity for studies of antibiotic resistance and evolutionary dynamics, which could contribute to improved management of patients with difficult-to-treat infections.

## Supplemental Material

Figure S1

Figure S2

Supplemental Materials and Methods

Supplemental Table 1

Supplemental Table 2

Supplemental Table 3

## Data Availability

The dataset generated during the current study are available in the Sequence Read Archive (SRA), BioProject PRJNA1330241.

## Acknowledgements

We gratefully acknowledge members of the UPMC Clinical Microbiology Laboratory and members of the Van Tyne laboratory for assistance throughout this study. This work was supported by the Cystic Fibrosis Foundation (grant SCON-00006650 to DVT), and by the Department of Medicine at the University of Pittsburgh (DVT). The funders had no role in study design, data collection and analysis, decision to publish, or preparation of the manuscript.

## References

1. WHO Bacterial Priority Pathogens List 2024: Bacterial Pathogens of Public Health Importance, to Guide Research, Development, and Strategies to Prevent and Control Antimicrobial Resistance. (World Health Organization, Geneva, 2024).

2. Letizia, M., Diggle, S. P. & Whiteley, M. Pseudomonas aeruginosa: ecology, evolution, pathogenesis and antimicrobial susceptibility. Nat. Rev. Microbiol. (2025) doi:10.1038/s41579-025-01193-8.

3. Shields, R. K. et al. Effectiveness of ceftazidime-avibactam versus ceftolozane-tazobactam for multidrug-resistant Pseudomonas aeruginosa infections in the USA (CACTUS): a multicentre, retrospective, observational study. Lancet Infect. Dis. 25, 574–584 (2025).

4. Pang, Z., Raudonis, R., Glick, B. R.Lin, T.-J. & Cheng, Z. Antibiotic resistance in Pseudomonas aeruginosa: mechanisms and alternative therapeutic strategies. Biotechnol. Adv. 37, 177–192 (2019).

5. Weimann, A. et al. Evolution and host-specific adaptation of Pseudomonas aeruginosa. Science 385, eadi0908 (2024).

6. Tam, V. H. et al. Prevalence, Resistance Mechanisms, and Susceptibility of Multidrug-Resistant Bloodstream Isolates of Pseudomonas aeruginosa. Antimicrob. Agents Chemother. 54, 1160–1164 (2010).

7. Bitar, I. et al. Genomic Characterization of Mutli-Drug Resistant Pseudomonas aeruginosa Clinical Isolates: Evaluation and Determination of Ceftolozane/Tazobactam Activity and Resistance Mechanisms. Front. Cell. Infect. Microbiol. 12, (2022).

8. Spottiswoode, N. et al. In host evolution of beta lactam resistance during active treatment for Pseudomonas aeruginosa bacteremia. Front. Cell. Infect. Microbiol. 13, (2023).

9. Raghuram, V. et al. Comparison of genomic diversity between single and pooled Staphylococcus aureus colonies isolated from human colonization cultures. Microb. Genomics 9, 001111 (2023).

10. Deurenberg, R. H. et al. Application of next generation sequencing in clinical microbiology and infection prevention. J. Biotechnol. 243, 16–24 (2017).

11. Chiu, C. Y. & Miller, S. A. Clinical metagenomics. Nat. Rev. Genet. 20, 341–355 (2019).

12. Cabot, G. et al. Evolution of Pseudomonas aeruginosa Antimicrobial Resistance and Fitness under Low and High Mutation Rates. Antimicrob. Agents Chemother. 60, 1767–1778 (2016).

13. Winstanley, C., O’Brien, S. & Brockhurst, M. A. Pseudomonas aeruginosa Evolutionary Adaptation and Diversification in Cystic Fibrosis Chronic Lung Infections. Trends Microbiol. 24, 327–337 (2016).

14. Abdelraouf, K., Kabbara, S., Ledesma, K. R., Poole, K. & Tam, V. H. Effect of multidrug resistance-conferring mutations on the fitness and virulence of Pseudomonas aeruginosa. J. Antimicrob. Chemother. 66, 1311–1317 (2011).

15. Liu, H. et al. Time Series Genomics of Pseudomonas aeruginosa Reveals the Emergence of a Hypermutator Phenotype and Within-Host Evolution in Clinical Inpatients. Microbiol. Spectr. 10, e0005722 (2022).

