## Supplementary figures and images for "Within-Patient Evolution of *Pseudomonas aeruginosa* Populations During Antimicrobial Treatment"

### Supplemental Fig. 1

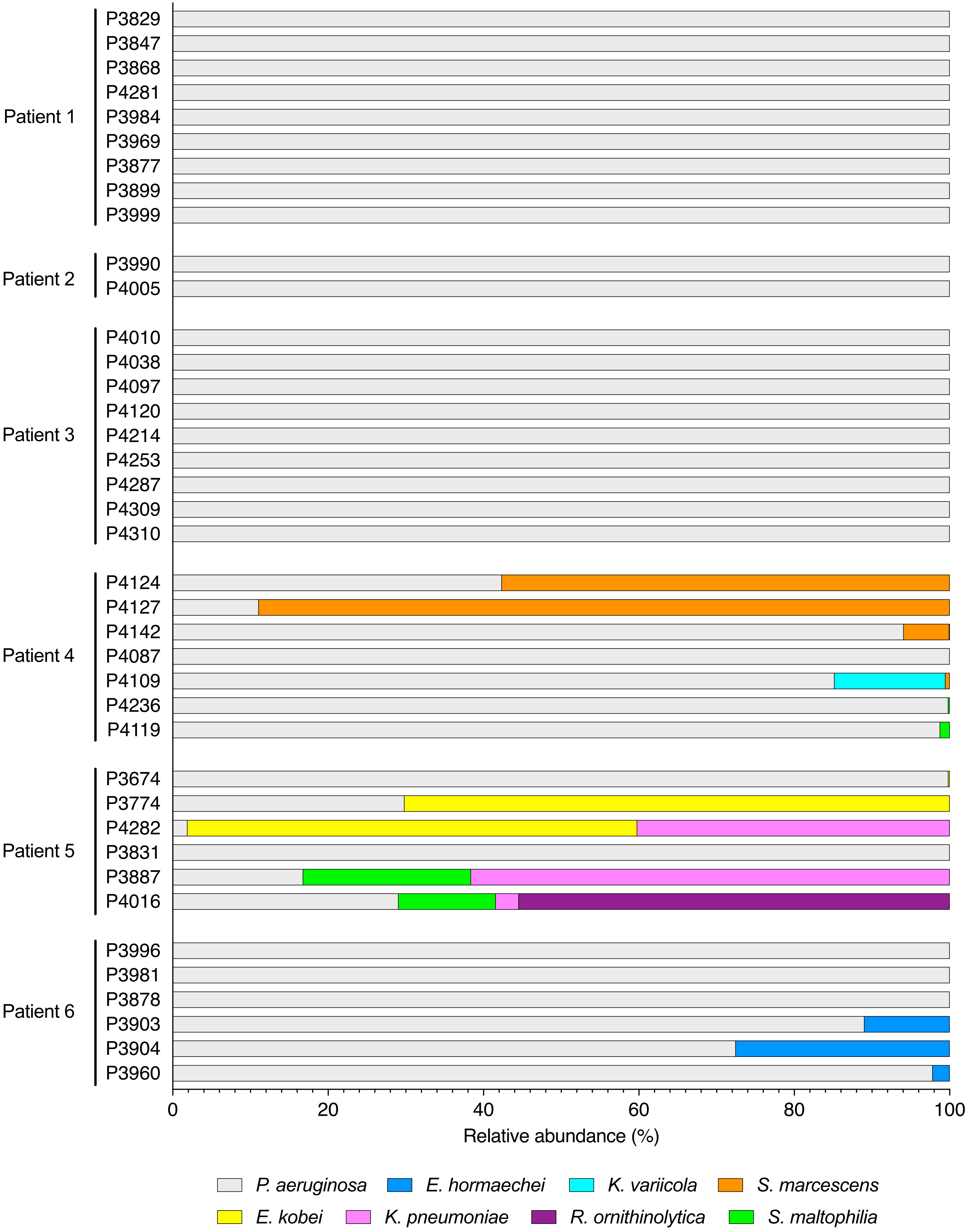

### Supplemental Fig. 2

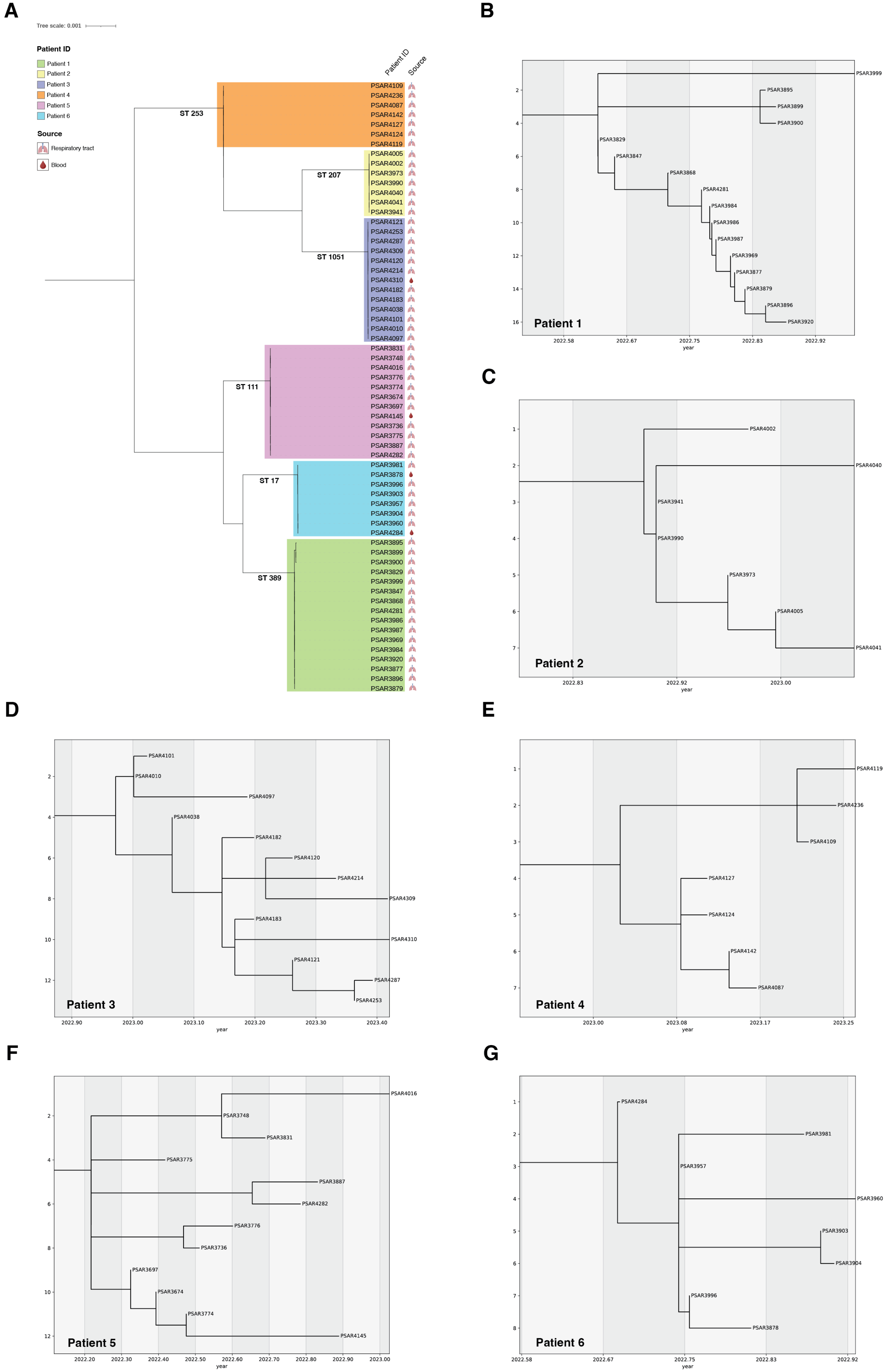
