## Supplemental File 1 for "Within-Patient Evolution of *Pseudomonas aeruginosa* Populations During Antimicrobial Treatment"

**Supplemental File 1. Patient Case Summaries**

**Patient 1.** 69-year-old male with history of COPD and idiopathic pulmonary fibrosis who presented for double lung transplant. Post transplantation, his hospital course was complicated by septic shock, respiratory failure requiring VV ECMO, sternal wound infection and dehiscence. He had prolonged respiratory failure and developed numerous hospital and ventilatory-acquired pneumonias due to *P. aeruginosa*. He ultimately died due to multiorgan failure in the hospital.

**Patient 2.** 85-year-old male with history of multiple myeloma, multiple falls resulting in intracranial hemorrhage, and cardiac arrest that was complicated by respiratory failure requiring prolonged mechanical ventilation. He developed recurrent *P. aeruginosa* ventilator-acquired pneumonia during a prolonged hospital stay. He eventually expired due to multiorgan failure and septic shock.

**Patient 3.** 57-year-old male with history of severe non-ischemic cardiomyopathy who was admitted for heart transplantation. His post operative course was complicated by cardiogenic shock, respiratory failure requiring mechanical ventilation, pressor support and VA ECMO. Throughout his critical illness, he had multiple infection-related complications including *P. aeruginosa* line-associated bacteremia and recurrent ventilator-associated pneumonia complicated by empyema. He eventually died due to chronic respiratory failure after a prolonged hospitalization.

**Patient 4.** 66-year-old male with history of nasopharyngeal squamous cell carcinoma status post-resection and chemotherapy, complicated by flap necrosis with fungal and polymicrobial bacterial infection requiring long-term antimicrobial treatment. He was re-admitted numerous times due to tumor progression, hemorrhagic complications, complex intra-abdominal infections, and septic shock from mixed sources. He had recurrent *P. aeruginosa* hospital-acquired pneumonia and was eventually discharged to a skilled nursing facility with palliative care.

**Patient 5**. 58-year-old male with history of hepatitis B, hepatitis C, and non-alcoholic fatty liver disease complicated by cirrhosis status post-deceased donor liver transplant. He developed colonic perforation due gastrointestinal mucormycosis requiring colectomy and ileostomy. During his hospitalization, he had respiratory failure and difficulty weaning from the ventilator, necessitating tracheostomy creation. He developed frequent *P. aeruginosa* hospital and ventilator-associated pneumonia throughout his multiple hospitalizations. The final hospitalization was complicated by septic shock and multiorgan failure, resulting in death.

**Patient 6.** 70-year-old male with history of alcohol use disorder complicated by chronic pancreatitis and metastatic prostate cancer who was admitted due sepsis secondary to cholangitis, bacteremia and fungemia. His course was complicated by respiratory failure requiring endotracheal ventilation. He had difficulty weaning from the ventilator, resulting in prolonged mechanical ventilation and tracheostomy. His course was further complicated by hemorrhage, ischemic colitis, and recurrent hospital and ventilator- associated pneumonias caused by *P. aeruginosa*. He died due to hemorrhagic shock resulting in multiorgan failure after a prolonged hospitalization.
