## Supplemental Methods for "Within-Patient Evolution of *Pseudomonas aeruginosa* Populations During Antimicrobial Treatment"

**Supplemental Materials and Methods**

**Sample Collection**

Longitudinal clinical specimens were collected from six patients infected with MDR *P. aeruginosa* who developed resistance to ceftolozane-tazobactam. For each patient, respiratory specimens were plated onto MacConkey agar, and unique *P. aeruginosa* morphologies were identified and reported as part of standard care in the clinical microbiology laboratory.

**Antimicrobial Susceptibility Confirmation**

Antibiotic susceptibility testing was performed using the MicroScan WalkAway system (Beckman Coulter, Brea, CA, USA), and resulted were interpreted according to the Clinical and Laboratory Standards Institute (CLSI) guidelines^1^. For study purposes, all colonies were scraped from the original MacConkey agar plate and pooled to generate culture-enriched metagenomic DNA samples for population-level analysis. This study was approved by the Institutional Review Board at the University of Pittsburgh under STUDY22070065. To confirm evolution of ceftolozane-tazobactam resistance for individual isolates, minimum inhibitory concentrations (MICs) of ceftazidime-avibactam (CZA) and ceftolozane-tazobactam (C/T) were determined by broth microdilution in triplicate according to standard CLSI methods as previously reported.

**Whole-Genome and Culture-Enriched Metagenomic Sequencing**

Isolates were then subjected to whole-genome sequence (WGS) analysis on the Illumina platform Genomic DNA was extracted using a Qiagen DNeasy Blood and Tissue Kit. Short-read libraries were prepared using the Illumina DNA Prep protocol and were sequenced (2 × 150 bp, paired-end) on a NextSeq 500. The earliest available isolate from each patient was used as a patient-specific reference, and a long-read library for each reference isolate was prepared using the Oxford Nanopore Technologies (ONT) rapid barcoding kit (SQK-RBK0004) and sequenced on the MinION platform.

**Data Analysis**

Assemblies were generated using SPAdes^2^ v3.15.5 (short reads) and unicycler^3^ v0.5.1 (short and long reads) and were assessed for quality using QUAST^4^. Species identification and contamination screening were performed with Kraken2^5^ v2.1.3. Multi-locus sequence typing (MLST) was carried out using the PubMLST database via *mlst*^6^ v2.11. Genome annotation was conducted with Prokka^7^ v1.14.5. Pairwise SNP distances and reference-free alignments of variants were computed using Split Kmer Analysis Toolkit (SKA) v1.0^8^. Resistance-associated mutations were identified from both single-colony and population-derived WGS data using *breseq*^9^. This study focused on a targeted set of 9 genes (Supplemental Table 1) known to be associated with antibiotic resistance. For culture-enriched metagenomic samples, *P. aeruginosa* genome-wide and individual gene coverage were calculated, and only samples with ≥100× genome-wide coverage were included in downstream analyses (Supplemental Table 1). Phylogenetic trees were constructed using RAxML HPC^10^ v8.2.12 with 100 bootstraps. Time-resolved phylogenies and ancestral sequence reconstruction were performed using TimeTree^11^ v0.11.4.

**References**

1.	Clinical and Laboratory Standards Institute (CLSI). Performance Standards for Antimicrobial Susceptibility Testing. in (CLSI, Wayne, PA, 2024).

2.	Bankevich, A. *et al.* SPAdes: a new genome assembly algorithm and its applications to single-cell sequencing. *J. Comput. Biol. J. Comput. Mol. Cell Biol.* **19**, 455–477 (2012).

3.	Wick, R. R., Judd, L. M., Gorrie, C. L. & Holt, K. E. Unicycler: Resolving bacterial genome assemblies from short and long sequencing reads. *PLOS Comput. Biol.* **13**, e1005595 (2017).

4.	Gurevich, A., Saveliev, V., Vyahhi, N. & Tesler, G. QUAST: quality assessment tool for genome assemblies. *Bioinformatics* **29**, 1072–1075 (2013).

5.	Wood, D. E., Lu, J. & Langmead, B. Improved metagenomic analysis with Kraken 2. *Genome Biol.* **20**, 257 (2019).

6.	Jolley, K. A. & Maiden, M. C. BIGSdb: Scalable analysis of bacterial genome variation at the population level. *BMC Bioinformatics* **11**, 595 (2010).

7.	Seemann, T. Prokka: rapid prokaryotic genome annotation. *Bioinformatics* **30**, 2068–2069 (2014).

8.	Harris, S. R. SKA: Split Kmer Analysis Toolkit for Bacterial Genomic Epidemiology. 453142 Preprint at https://doi.org/10.1101/453142 (2018).

9.	Deatherage, D. E. & Barrick, J. E. Identification of mutations in laboratory-evolved microbes from next-generation sequencing data using breseq. *Methods Mol. Biol. Clifton NJ* **1151**, 165–188 (2014).

10.	Stamatakis, A. RAxML version 8: a tool for phylogenetic analysis and post-analysis of large phylogenies. *Bioinformatics* **30**, 1312–1313 (2014).

11.	Kumar, S., Stecher, G., Suleski, M. & Hedges, S. B. TimeTree: A Resource for Timelines, Timetrees, and Divergence Times. *Mol. Biol. Evol.* **34**, 1812–1819 (2017).
